# Function of the bacteriophage P2 baseplate central spike Apex domain in the infection process

**DOI:** 10.1101/2023.02.25.529910

**Authors:** John-Mark Miller, Ekaterina S. Knyazhanskaya, Sergii A. Buth, Nikolai S. Prokhorov, Petr G. Leiman

**Author notes:** Emory University, School Of Medicine, Department of Pediatrics, Infectious Diseases, 2015 Uppergate Dr., Emory Children’s Center Bldg., Atlanta, Georgia 30322 (Sergii A. Buth).

## Abstract

The contractile tail of bacteriophage P2 functions to drive the tail tube across the outer membrane of its host bacterium, a prerequisite event for subsequent translocation of phage genomic DNA into the host cell. The tube is equipped with a spike-shaped protein (product of P2 gene *V*, gpV or Spike) that contains a membrane-attacking Apex domain carrying a centrally positioned Fe ion. The ion is enclosed in a histidine cage that is formed by three symmetry-related copies of a conserved HxH (histidine, any residue, histidine) sequence motif. Here, we used solution biophysics and X-ray crystallography to characterize the structure and properties of Spike mutants in which the Apex domain was either deleted or its histidine cage was either destroyed or replaced with a hydrophobic core. We found that the Apex domain is not required for the folding of full-length gpV or its middle intertwined β-helical domain. Furthermore, despite its high conservation, the Apex domain is dispensable for infection in laboratory conditions. Collectively, our results show that the diameter of the Spike but not the nature of its Apex domain determines the efficiency of infection, which further strengthens the earlier hypothesis of a drill bit-like function of the Spike in host envelope disruption.

## Introduction

The tail of bacteriophage P2 (**Fig. 1a, 1b**) is an archetypical representative of Contractile Injection Systems (CISs) – macromolecular machines that use a contractile sheath and rigid tube to breach the envelope of a target cell [1, 2]. In all known CISs, the membrane-attacking end of the tube carries a spike-shaped protein aka Baseplate Central Spike [3-16] (simply Spike here for brevity, **Fig. 1a**).

**Figure 1.**
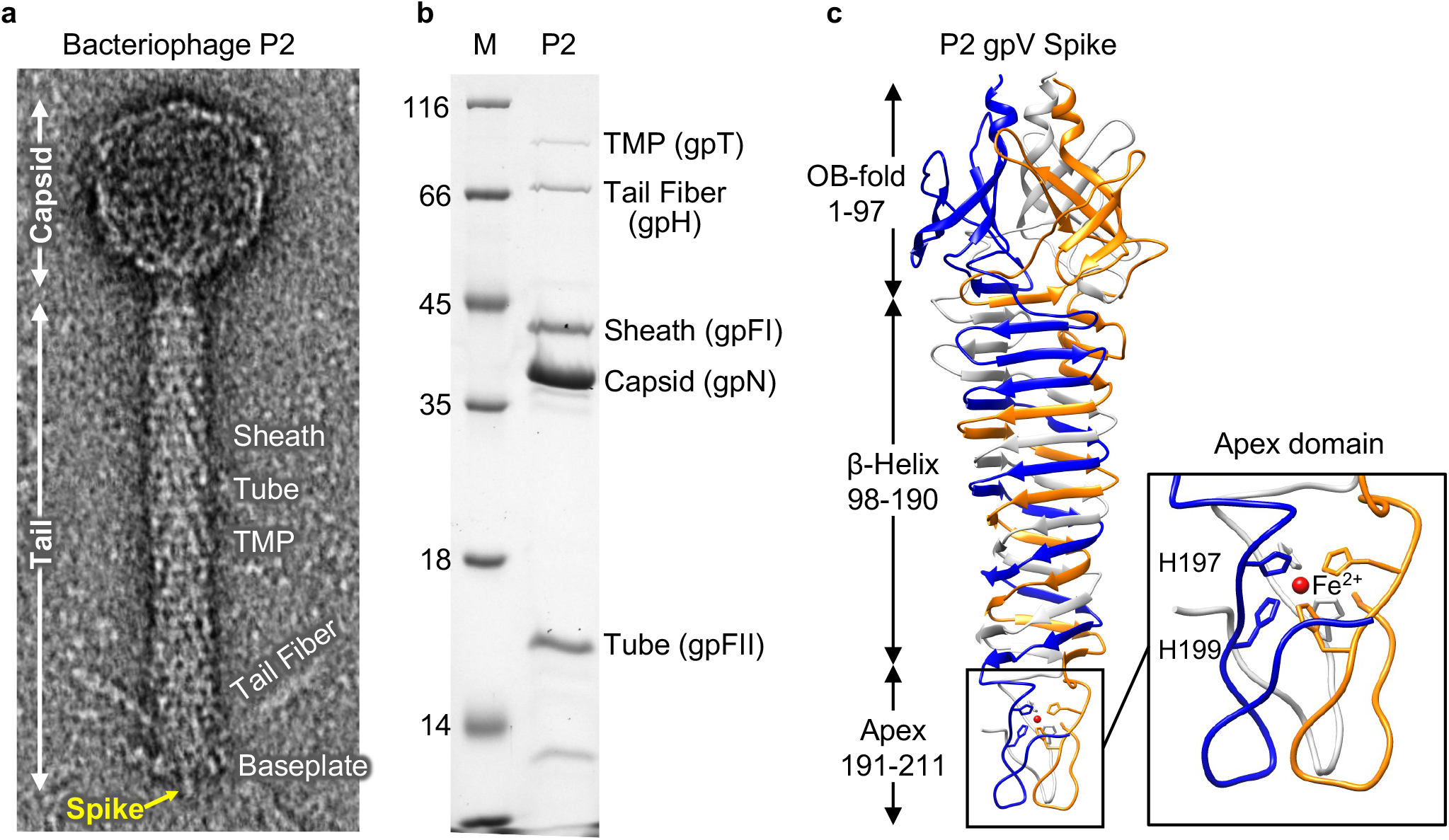
Bacteriophage P2 and its baseplate Spike. **a**, Electron micrograph of negatively stained (by uranyl acetate) phage P2 vir1 particle. The main components of the particle are labeled. **b**, Coomassie-stained sodium dodecyl sulfate gel electrophoresis (SDS-PAGE) of CsCl-gradient purified bacteriophage P2 vir1. Left lane -molecular weight marker. The numbers correspond to molecular weights in kDa. **c**, Ribbon diagram of P2 gpV. Each chain is colored in a distinct color. Three copies of histidines H197 and H199 form an octahedral intermolecular histidine cage, which traps an iron ion (colored red).

Bacteriophage P2 Spike (the product of gene *V*, gpV) consists of an N-terminal oligonucleotide/oligosaccharide-binding fold (OB-fold) domain, which attaches the Spike to the tube/baseplate, an intertwined triple-stranded β-helical domain, and a conical Apex domain [3, 17] (**Fig. 1c**). A characteristic and universally conserved feature of the Apex domain is a buried, centrally positioned Fe^2+^ ion, which is coordinated by six histidine sidechains in an octahedral configuration [3] (**Fig. 1c**). This intermolecular histidine cage is formed by three symmetry-related copies of the HxH motif (histidine, any residue, histidine) (**Fig, 1c**). The HxH motif is the only universally conserved sequence feature of Spikes in unrelated contractile tail phages (**Fig. 2**). As a consequence, the presence of the HxH motif in the C-terminal part of a Spike protein sequence can be used to identify an Apex domain [3, 4]. Spike proteins whose sequence does not contain an HxH motif near the C-terminus (e.g. phage T4 gp5) carry a PAAR (proline-alanine-alanine-arginine) repeat protein (e.g. T4 gp5.4), which could be homologous to the Apex domain [13].

**Figure 2.**
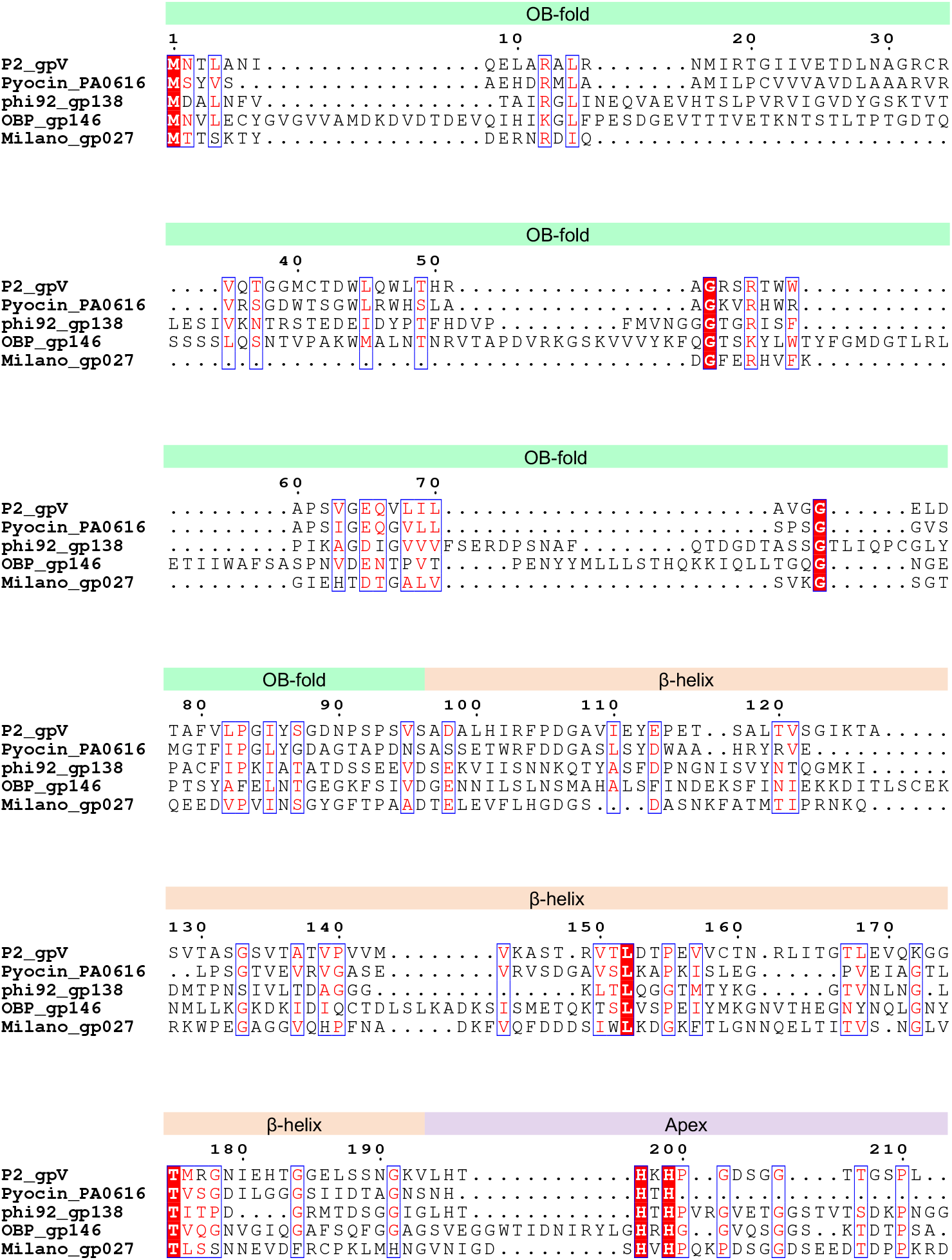
The HxH histidine cage motif is conserved in Spikes from diverse contractile tail bacteriophages and tailocins. Sequences of *Escherichia coli* phage P2 gpV, *Pseudomonas aeruginosa* PAO1 R-type pyocin gene product PA0616, *E. coli* phage phi92 gp138, *P. fluorescens* jumbophage OBP gp146 and *Agrobacterium tumefaciens* phage Milano gp027 aligned using MAAFT [50] using default parameters and visualized using ESPript [51]. The residues are numbered as in P2 gpV.

Besides Spikes, the intermolecular HxH histidine cage motif is found in phage tail fibers and tailspikes where it is thought to stabilize the trimeric fold and/or function during folding by keeping the polypeptide chains in register [18-21]. These hypotheses have not been tested experimentally. The roles of the Fe^2+^ ion and the histidine cage of the Spike in phage infection are also unknown.

Here, we used solution biophysics and X-ray crystallography to characterize the structure and properties of three mutants of the bacteriophage P2 gpV Spike in which the Apex domain was either deleted or both of its Fe^2+^ ion-binding histidines were replaced with either alanines or phenylalanines. Furthermore, we introduced these mutations to the bacteriophage P2 genome and quantified the impact of altering the

Apex domain structure on the efficiency of phage infection. Lastly, we analyzed the impact of these mutations on the cellular localization of the Spike upon infection. Our findings reveal the role of the histidine cage motif in the folding of the Apex domain and in phage infection.

## Results

### GpV mutants and outline of work

The Apex domain of gpV is formed by residues 191-211, and the Fe-coordinating histidine cage – by the side chains of His197 and His199 (**Fig. 1c**). We reasoned that the Fe-binding site can be destroyed by replacing both histidines with alanines. To test whether a hydrophobic core can function in place of the histidine cage, we replaced both histidines with phenylalanines. We called the H197A/H199A and H197F/H199F mutants gpV-AA (or AA) and gpV-FF (or FF), respectively. We also created a mutant in which the Alex domain was deleted – gpV-DEL (or DEL). The cloning procedure inadvertently introduced an exogenous leucine (Leu197) downstream from the last native residue Thr196.

In the *in vitro* part of the work, biochemical, biophysical, and structural properties of the three gpV mutants have been compared to each other and to those of the wild-type (WT) gpV. In the *in vivo* part, the gpV mutations and WT gpV were introduced into the P2 vir1 phage genome (strictly lytic P2 mutant [22]) by way of a *de novo* constructed intermediate P2 vir1 *Vam1* mutant. The infection properties of these mutant phages were then compared to each other. The cellular localization of WT gpV and its Apex domain mutants from the infecting phage particle has been established by cell fractionation and immunoblotting.

### Solution biophysics studies of P2 gpV mutants

The WT and mutant gpV proteins were expressed in *Escherichia coli* (*E. coli*) BL21 (DE3) and purified to homogeneity (**Fig. 3a, 3b**). All three mutants, including DEL, were soluble showing that the Apex domain is dispensable for folding or trimerization of the protein. In size exclusion chromatography (**Fig. 3a**), the AA and FF mutants eluted earlier than the WT and DEL indicating that the former have a larger radius of gyration possibly because of disorder in their Apex domains.

**Figure 3.**
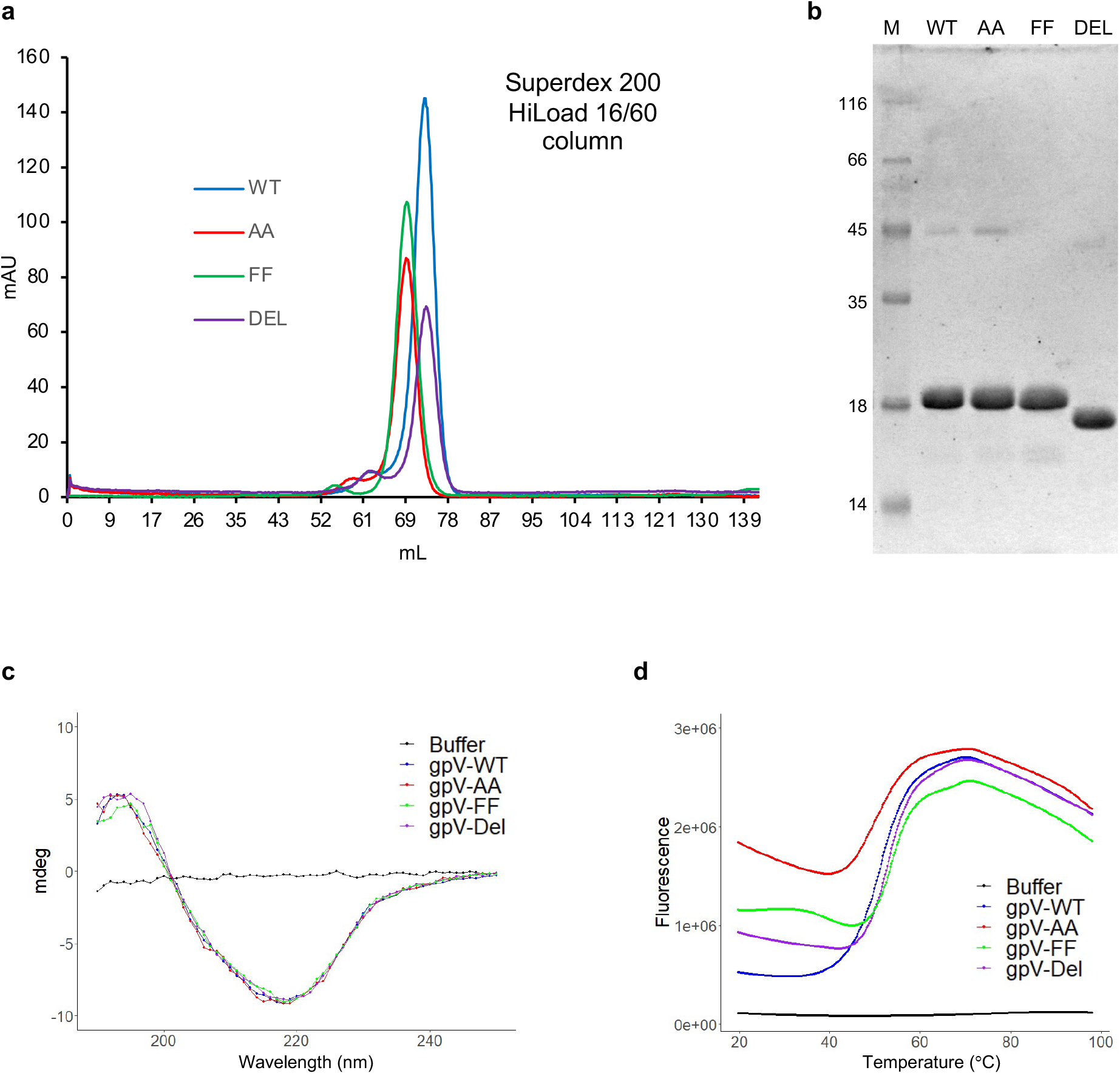
Solution biophysics and biochemical characterization of gpV with altered Apex domain. **a** and **b**, Size exclusion chromatography and 12% sodium dodecyl sulfate gel electrophoresis, respectively, of WT gpV and the three mutants. **c** and **d**, CD and DSF of WT gpV and the three mutants. The CD curves are averages of two biological replicates with three technical replicates each. The DSF curves are averages of two biological replicates with six technical replicates each.

To gain deeper insight into the structure of gpV mutants, we performed circular dichroism (CD) spectroscopy of WT gpV and its three mutants (**Fig. 3c**). In the β-helical domain, the secondary, tertiary, and quaternary structures are inseparable, so any alteration to the secondary structure, which could be measured in CD, would indicate that the tertiary and quaternary structures are also affected.

The CD spectra of the WT and mutant samples were virtually identical with a positive peak around 195 nm and a negative peak at 218 nm, as is seen in other predominately β-structural proteins (**Fig. 3c**). These results showed that altering the histidine cage structure or deleting the Apex domain had minimal to no impact on the structure of the β-helical domain. Furthermore, the Apex domain appeared to contribute little to the CD spectra of the full-length protein.

To examine the level of disorder in mutants with altered histidine cages, we performed differential scanning fluorimetry (DSF) [23]. The protein samples were mixed with a hydrophobic dye SYPRO orange and heated in the interval of 20-99°C (**Fig. 3d**). The DSF curve of WT gpV was virtually flat in the interval of 20-40°C showing that WT gpV is a well-folded protein with few hydrophobic regions exposed to solution. The DEL mutant displayed a greater signal in the same interval, showing that the SYPRO dye partially penetrated into or interacted with the hydrophobic interior of the β-helical domain through its C-terminal solution-exposed end. In the AA mutant, the amount of fluorescence was about four times greater than that in the WT suggesting that the Apex domain was fully disordered and all of its hydrophobic elements were accessible to the dye. In the FF mutant, the fluorescence signal was significantly lower than in the AA mutant, despite the addition of two large hydrophobic side chains. This suggests that in the FF mutant, the Apex domain was partially or possibly nearly fully ordered. All fluorescence curves display a downward trend with increasing temperature because the mobility of water molecules, the main source of SYPRO orange fluorescence quenching, increases with temperature.

Notably, the first inflection point in all DSF spectra, which is often considered to be the temperature of unfolding (or melting temperature), was very similar for all variants (**Fig. 3d**) and was located between 50°C and 55°C. This is much lower than 73° C measured using differential scanning calorimetry for the orthologous and structurally similar Spike of phage Mu [16]. It is also incompatible with the previously measured half-life of the P2 gpV trimer, which was 37 min at 65°C as visualized using SDS-PAGE [3]. Thus, the first peak in the DSF curve likely corresponded to the denaturation of the N-terminal OF-fold domain. Nevertheless, the exact temperature of denaturation is of no importance in this assay, as it was set up to evaluate the amount of disorder in protein structure at room temperature.

### Crystal structure of gpV-AA, gpV-FF, and gpV-DEL

The solution biophysics experiments and size exclusion chromatography described above showed that 1) the Apex domain plays no role in folding or stability of the Spike; 2) the Apex domain could be partially folded in the FF mutant; and 3) the Apex domain was likely completely disordered in the AA mutant. To strengthen these experimental findings, we determined the structure of these mutants by X-ray crystallography. As the Apex domain is not well-defined in the X-ray map of full-length gpV [3], we transferred these mutations to a shorter gpV mutant (residues 97-211, β-helix plus Apex), previously called gpVdL (**Fig. 4a**), which was known to crystallize easily and be fully ordered [10].

**Figure 4.**
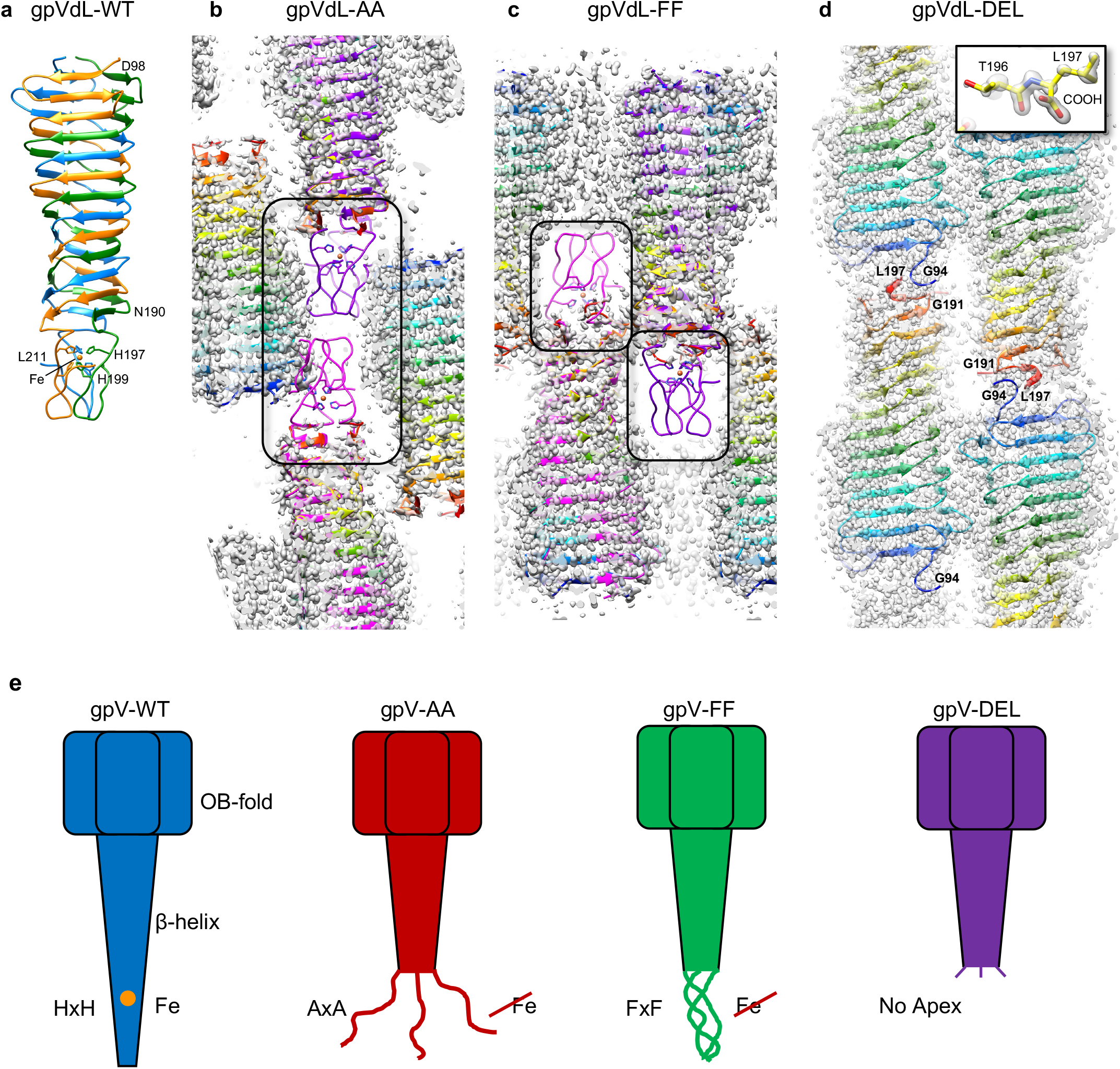
X-ray crystallographic analysis of gpVdL Apex mutants. **a**, Ribbon diagram of the crystal structure of WT gpVdL. **b-d**, Electron density maps (contoured at 1.5 standard deviations above the mean) of gpVdL-AA, gpVdL-FF and gpVdL-DEL crystals demonstrating the ordered and disordered regions of atomic models. The atomic models of gpVdL-AA, gpVdL-FF and gpVdL-DEL are shown in cartoon/ribbon representation and are colored in visibile spectrum colors from blue (N-terminus) to red (C-terminus). The atomic models of two gpVdL WT trimers (colored magenta and purple and also in ribbon/cartoon representation) are superimposed onto two gpVdL-AA and gpVdL-FF trimers comprising the asymmetric unit. The Apex domains are boxed in black rectangles. The inset in panel **d** shows the atomic model and the electron density of the C-terminal residues for one of the chains of the gpVdL-DEL mutant. **e**, Schematics showing the structure of the Apex domain in the WT and three mutants based on all biochemical, solution biophysics, and structural data.

The gpVdL-AA, gpVdL-FF, and gpVdL-DEL mutants crystallized in conditions close to those of the WT and diffracted X-rays to very high resolution (**Table 1**). While the electron density of the β-helix is well defined in all three mutants, the density corresponding to the Apex domain is absent in both the AA and FF mutant crystals (**Fig. 4b, 4c**). Superposition of the WT gpVdL structure onto the ordered part of the gpVdL-AA mutant shows that a folded Apex domain would clash with other molecules in the crystal lattice (**Fig. 4b**). To the contrary, the crystal lattice of the FF mutant can accommodate a folded Apex domain (**Fig. 4c**). The very C-terminal element of the DEL mutant (residues 191-197), which correspond to the N-terminal part of the Apex domain, does not display threefold symmetry and its electron density is worse than that of the rest of the protein (**Fig. 4d**). Nevertheless, the atomic model of the DEL mutant can be built to the very C-terminal residue (**Fig. 4d, inset**).

**Table 1.**
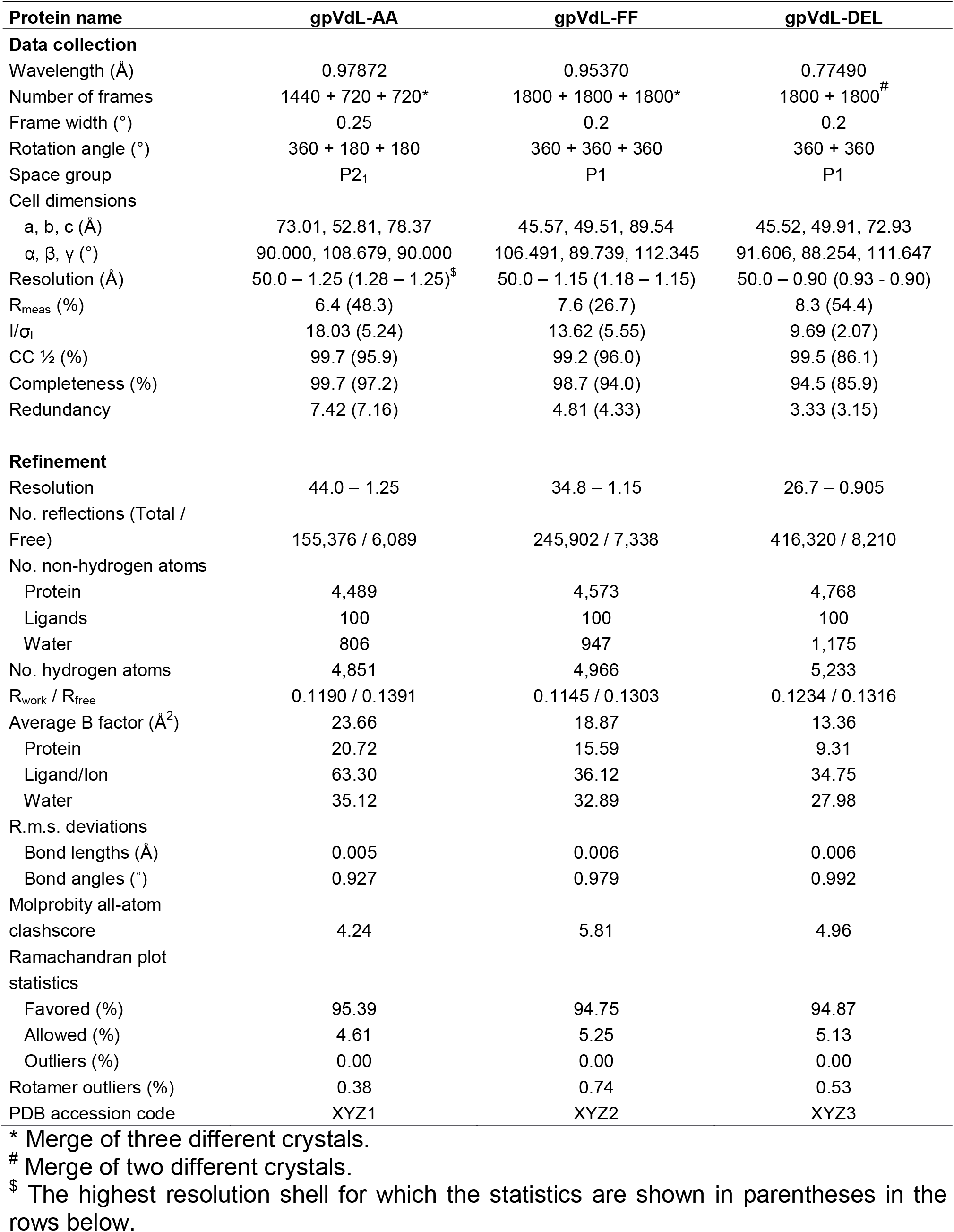
Crystallographic statistics.

Thus, the X-ray diffraction data demonstrate excellent agreement with the findings of solution biophysics experiments and provide additional detail. In solution and *in crystallo*, the Apex domain of the gpV-AA mutant is fully disordered. The FF mutant’s Apex is likely folded *in crystallo* but is positionally disordered, similar to full-length gpV where the X-ray map of the Apex domain was too poor for atomistic interpretation [3]. Of note, the absence of electron and cryo-EM density of fully folder domains is a common occurrence in other proteins (see e.g. [24]). In the DEL mutant, the C-terminal element of the protein exhibits some mobility compared to the β-helix (**Fig. 4e**).

### The sharpness of gpV Spike impacts phage P2 infectivity

To examine the role of the Apex domain and the delivery destination of gpV during phage P2 infection, the Apex domain mutations described above had to be introduced into the phage genome. To do so, we first created a P2 vir1 *Vam1* mutant phage, which carried an amber stop codon in place of the ninth amino acid of gene *V* (Gln9-Am) (**Fig. 5a-5d**). P2 vir1 *Vam1* could be propagated on the amber-suppressing *E. coli* strain C-520 (*supD*, UAG-tRNA^ser^) but did not grow on non-suppressing *E. coli* C-2 (laboratory host of P2 vir1) [25, 26] (**Fig. 5d**). Then, we replaced the *Vam* gene of P2 vir1 *Vam1* with the WT gene *V* and with gene *V* carrying Apex domain mutations described above (**Fig. 5e-5h**). These mutant phages – P2 vir1 *Vam1-WT*, P2 vir1 *Vam1-AA*, P2 vir1 *Vam1-FF* and P2 vir1 *Vam1-DEL* – grew on non-suppressing *E. coli* C-2. For both, sense-to-nonsense and nonsense-to-sense phage genome mutagenesis, we used the phage λ red recombinase system [27, 28].

**Figure 5.**
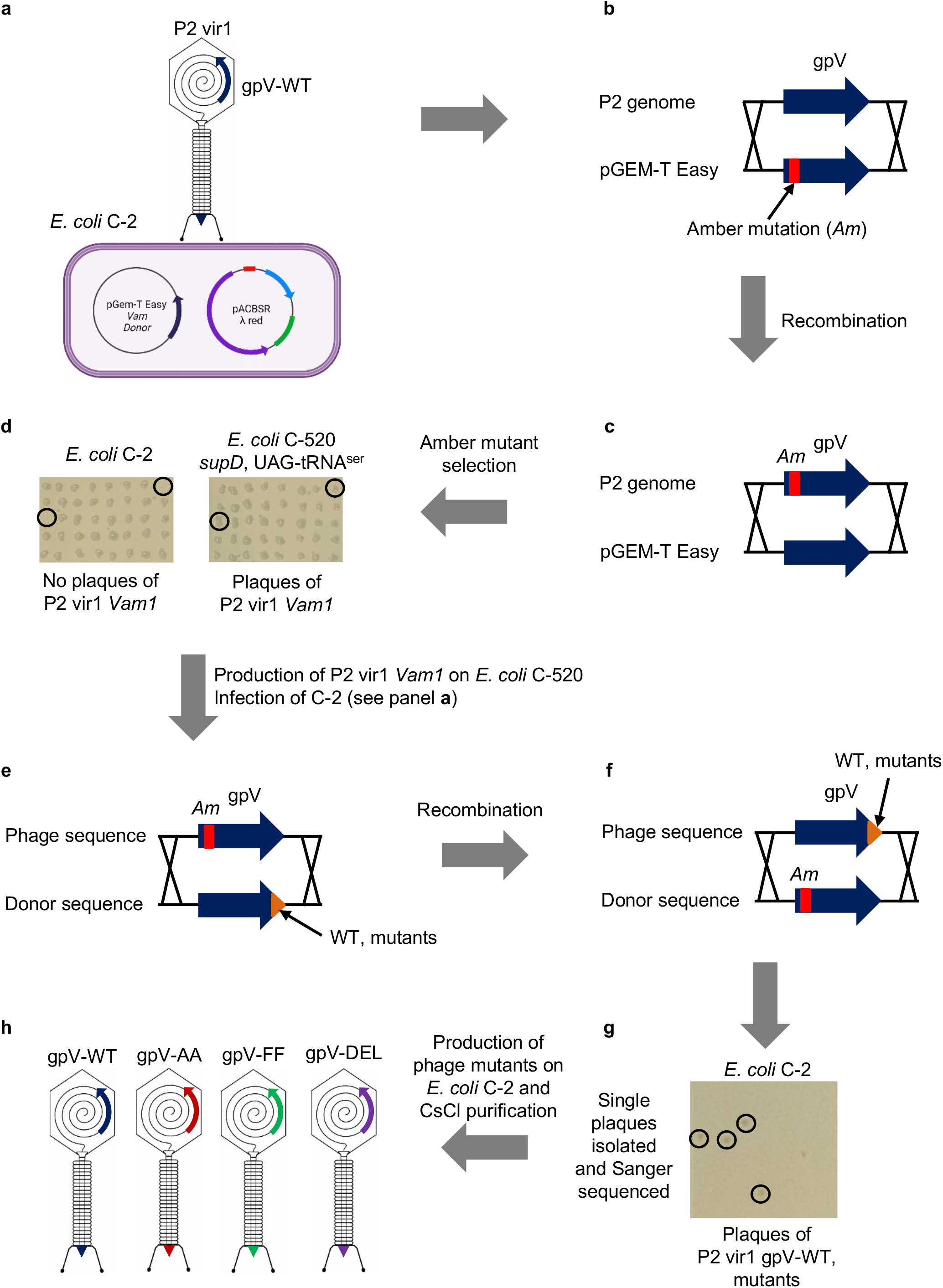
Flowchart outlining phage mutagenesis. **a**, Plasmids used for recombination during phage infection. pGem-T Easy (Amp^r^) contained a fragment of P2 genome encompassing gene *V* carrying a nonsense (amber) mutation at codon 9 (Gln9-Am). pACBSR (Cam^r^) contained the phage λ red recombination system genes. The cell schematic was created with BioRender.com. **b, c**, The recombination event. **d**, Selection of P2 vir1 *Vam1* with the help of non-permissive (C2) and permissive (C-520) *E. coli* strains. **e, f**, Recombination to restore the WT sequence of gene V and to introduce mutations into the Apex domain. **g, h**, Selection of P2 vir1 mutants carrying WT and mutated Apex domains.

For quantification of infection efficiency, P2 vir1 *Vam1*-WT/AA/FF/DEL mutants were first purified using a CsCl density gradient (**Fig. 6a**). This procedure separates the phage, a composite DNA-protein particle with a unique buoyant density, from nucleic acid, lipid, and protein impurities. However, the buoyant density of tailless, DNA-packaged capsids is close to that of mature, fully assembled phages, and the two can co-purify in CsCl gradient ultracentrifugation. To account for this possibility, the count of mature, fully assembled phages in all mutants was derived from the amount of tail sheath (gpFI) and tail tube (gpFII) proteins present in the sample. Namely, the titers of all phage preparations were normalized to the intensities of their gpFI and gpFII bands in Coomassie-stained SDS-PAGE gels (**Fig. 6a**).

**Figure 6.**
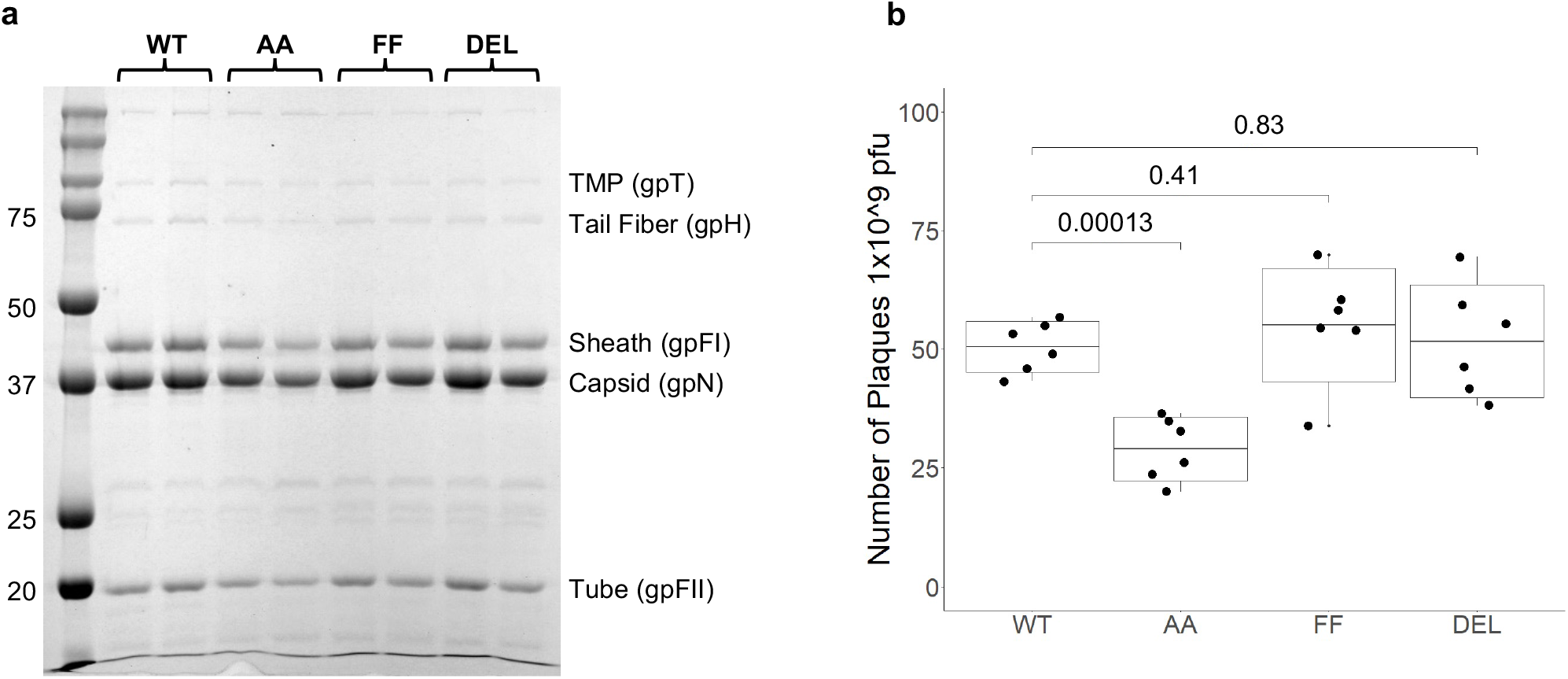
The role of Apex domain in phage infection as determined by the EOP assay. **a**, SDS-PAGE of CsCl-purified stocks of P2 vir1 mutants carrying the WT Spike, a Spike with a disordered Apex domain (the AA mutant), a Spike with a partially folded Apex (the FF mutant) and a Spike without Apex (the DEL mutant). Two biological replicates for each mutant are shown. **b**, Boxplot of EOPs measured by a standard double layer agar assay of the WT and three mutants shown in panel **a**. The p-values are calculated using an unpaired Student’s t-test.

The efficiency of plating (EOP) or the number of infection-capable particles per unit volume of P2 vir1 *Vam1*-WT/AA/FF/DEL mutants was quantified using a standard double-layer agar plaque assay. P2 vir1 *Vam1-WT*, P2 vir1 *Vam1-FF*, and P2 vir1 *Vam1-DEL* demonstrated similar EOPs, whereas the EOP of P2 vir1 *Vam1-AA* was about twofold lower (**Fig. 6b**). As the EOPs of P2 vir1 *Vam1-WT* and P2 vir1 *Vam1-DEL* were very similar, these experiments also showed that the Apex domain of the Spike has no function in phage assembly.

### WT gpV and its Apex domain mutants are translocated to the periplasm upon membrane penetration

In the course of a normal infection process, sheath contraction results in the delivery of the WT P2 gpV Spike into the periplasm of the host cell (Mattenberger et al.). However, mutations in the gpV Apex domain could alter the final location of gpV and hinder the infection process. The mutated gpV could become stuck in the outer membrane affecting subsequent processes such as, for example, the exit of the tape measure protein from the tail tube.

The location of the WT gpV Spike and its Apex domain mutants in the infected cell was established with the help of a Western blot-based assay similar to the one described in Mattenberger et al. with a slight modification (**Fig. 7a**). Unlike P2 vir1 used in the Mattenberger et al. study, P2 vir1 *Vam1*-WT/AA/FF/DEL mutants could grow on non-permissive *E. coli* C-2. To prevent transcription of phage genes and synthesis of new copies of gpV-WT/AA/FF/DEL that would interfere with the results, the infection was performed in the presence of rifampicin [29]. The latter did not interfere with phage adsorption as all mutants demonstrated similar adsorption efficiencies. This finding also showed that the Apex domain does not play a role in the adsorption process (**Fig 7b**).

**Figure 7.**
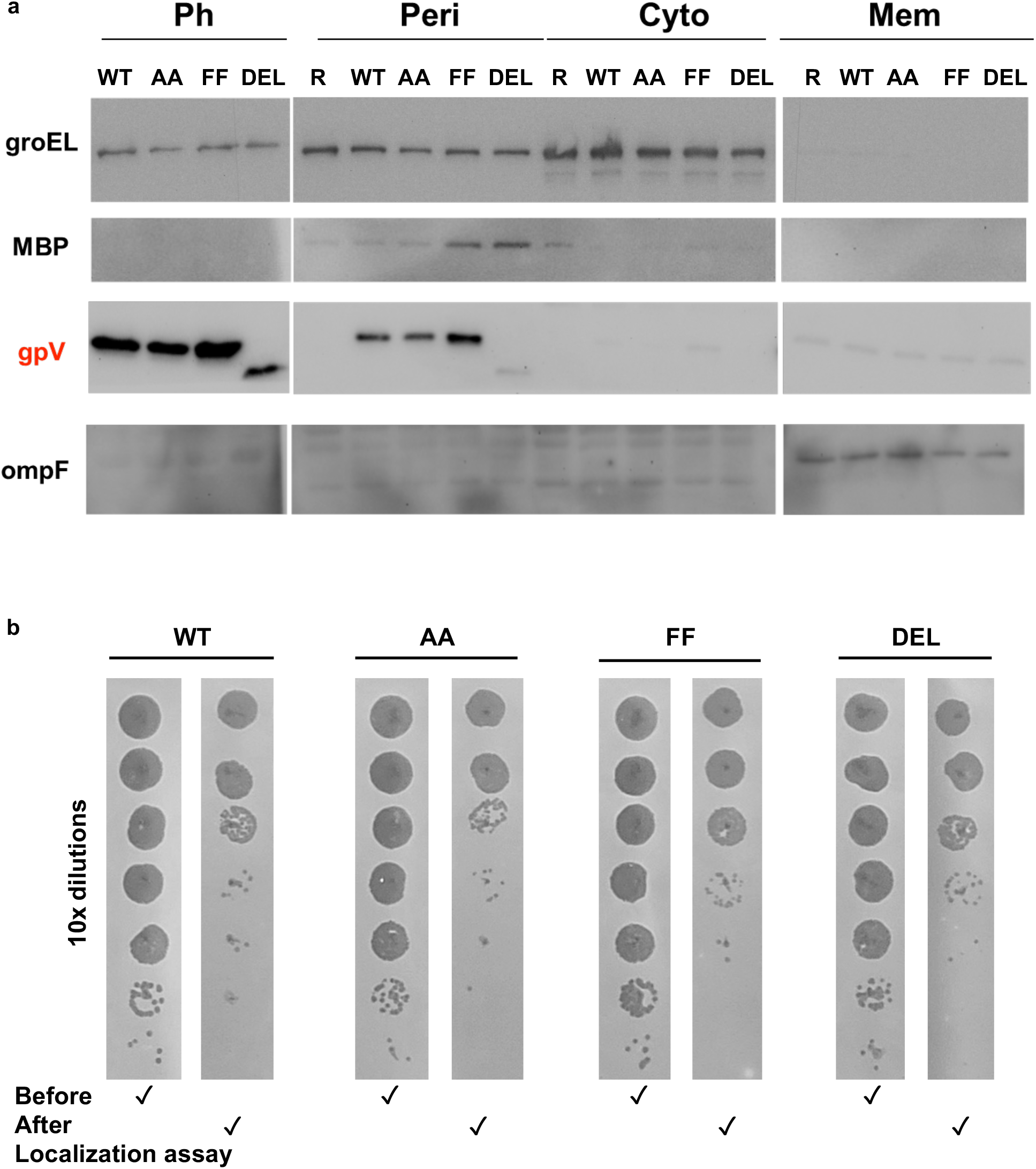
Cellular localization of WT gpV and its Apex domain mutants originating from the infecting virus particle. **a**, Phages (WT and the three mutants) and three cellular fractions – the periplasm (Peri), the cytoplasm (Cyto) and the membranes (Mem) – were blotted with antibodies against cellular markers (groEL for cytoplasm, MBP (maltose-binding protein) for periplasm, and ompF for membranes) and with anti-gpV antibodies. R stands for rifampicin-treated uninfected cells (the negative control). The immunogenicity of gpV-DEL, in which the Apex domain is deleted, is poor. The cellular marker antibodies are monoclonal whereas the anti-gpV anibodies are polyclonal. **b**, Adsorption assay of P2 vir1 carrying WT gpV and its Apex domain mutants. For each mutant, the top and bottom panels show 10-fold titration of the phage stock prior and after its use in the gpV localization assay, respectively.

To confirm that the fractionation procedure divided the infected cell lysates into correct fractions, we assayed them for the presence of proteins with known cellular localizations with the help of appropriate antibodies. The maltose binding protein (MBP), GroEL, and OmpF (outer membrane protein F) served as markers for the periplasmic, cytoplasmic, and combined inner and outer membrane fractions, respectively (**Fig. 7a**).

The localization assay showed that, similar to the WT gpV, all three mutants localized to the periplasm (**Fig. 7a**). GpV-DEL reacted poorly with our custom polyclonal antibodies, which were used in all Western blot assays and which were raised against recombinant full length gpV. This shows that the Apex domain is important for gpV immunogenic properties.

## Discussion

Bacteriophage-encoded homologous proteins demonstrate some of the highest amino acid sequence divergence [24, 30, 31]. In particular, phage virion proteins, while being homologous and structurally similar, have markedly different amino acid sequences [2, 3, 12, 32, 33]. Position-dependent conservation of sequence features in structural elements of bacteriophage virion proteins is extremely rear and is often associated with enzymatic activity [34]. To this end, Apex domains of Spike proteins demonstrate exceptionally high conservation of the HxH cage, which is akin to the conservation of catalytic site residues in enzymes (**Fig. 2**). This observation prompted us to investigate the role of the histidine cage in the folding of the phage P2 Spike, its Apex domain, and the function of the cage in phage infection.

We found that the Apex domain does not play a substantial role in the folding of the rest of the Spike – that is the β-helical domain folds independently of the Apex (**Fig. 3**), which mirrors findings for the phage Mu central spike [16]. Destruction of the histidine cage (as in the gpV-AA mutant) caused the Apex domain to unfold but the structure of the rest of the protein appears to be unaffected (**Fig. 3c, 3d, 4b, 4e**). Replacement of the histidine cage with a hydrophobic core of a similar size (the gpV-FF mutant) resulted in a domain, which is either in a constant transition between a folded and unfolded state or is folded but disordered positionally (**Fig. 3c, 3d, 4c, 4e**).

Rather unexpectedly, we found that the histidine cage and the Apex domain are dispensable for infection in laboratory conditions. Deletion of the Apex domain results in a phage mutant whose infection properties are indistinguishable from that carrying WT gpV (**Fig. 6b**). Furthermore, replacement of the histidine cage with a hydrophobic core also results in a WT-like phenotype (**Fig. 6b**), which further indicates that in this mutant the Apex domain is folded. The EOP or the infection efficiency of the AA phage mutant carrying a fully unfolded Apex and thus having an effectively larger diameter spike is about two times lower than that of the WT (**Fig. 6b**).

Our work also makes it possible to examine whether machine learning-based protein structure prediction approaches such as AlphaFold [35] and RoseTTAFold [36] can give insight into the effect of histidine cage mutations on the structure of the Apex domain. AlphaFold models of AA and FF mutant Apex domains are virtually identical to that of the WT (**Fig. 8**). Surprisingly, the confidence of the most disordered AA mutant Apex model is highest, essentially experimental level-like (**Fig. 8a**). The structure of the FF mutant Apex is predicted with a lower, albeit still significant, confidence (**Fig. 8b**), so without experimental data, one could hypothesize that this Apex domain is less ordered than the AA Apex. This clearly disagrees with all experimental data presented in this paper, which demonstrates that machine learning-based structure prediction is unsuitable for the analysis of critical point mutations.

**Figure 8.**
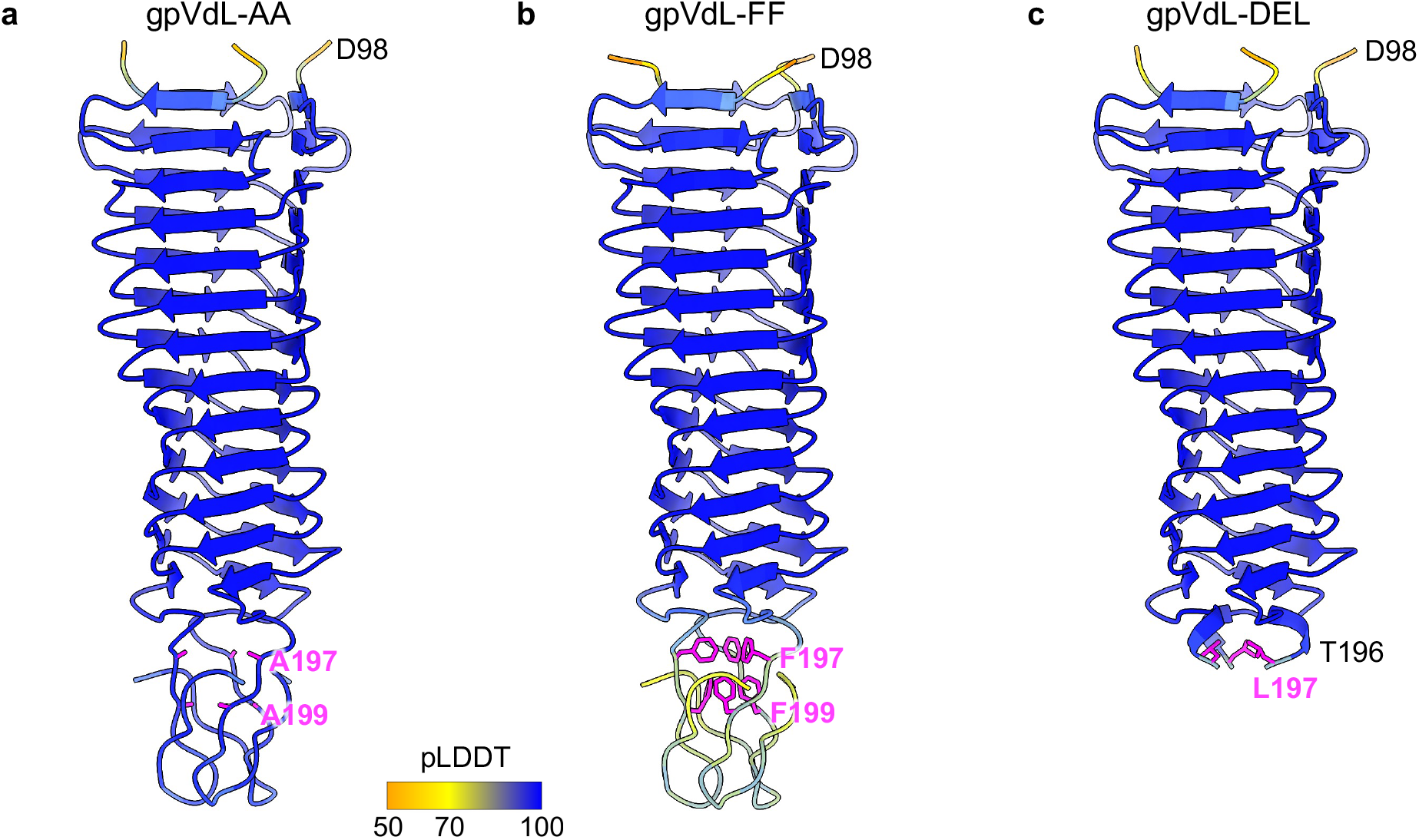
AlphaFold models of gpVdL Apex domain mutants. **a, b**, and **c**, Ribbon diagrams of AlphaFold models of gpVdL-AA, gpVdL-FF and gpVdL-DEL mutants colored according to their confidence, which is depicted with the help of a color bar that is shown below panels **a** and **b**. pLDDT stands for predicted local distance difference test in which a higher value represents higher model confidences with a value of 100 corresponding to a model that is indistinguishable from experimental data [35]. The side chains of mutated residues are colored magenta.

Summarizing all results, we conclude that only the cross-sectional area of the membrane-attacking tip of the Spike is important for phage P2 infectivity in laboratory conditions. This finding supports an earlier hypothesis that Spikes act as mechanical drill bits in contractile phage tails and tailocins [3]. Our experiments show that the Apex domain with a hydrophobic core may be partially unfolded and thus is “softer” than the WT variant with its Fe-carrying histidine cage. Even though this “softness” does not result in a phenotype in laboratory conditions, extremely high conservation of the histidine cage in phage baseplate central Spike proteins suggests that it is likely to be important for infection in multispecies environments containing bacterial hosts carrying diverse lipopolysaccharide structures.

## Materials and Methods

### Mutagenesis and cloning

Plasmids containing P2 phage full-length gpV and N-terminal truncation mutant gpVdL were constructed as previously described [3]. These plasmids were mutated to contain either one of the double substitution mutations or the deletion mutant by overlapping PCR [37] using primers listed in **Supplementary Table 1**. PCR was conducted using CloneAmp HiFi PCR Premix (Clonetech) and the following thermocycler conditions: 1) initial denaturation at 99°C for 30 seconds, 2) 35 cycles of denaturation at 99°C for 10 seconds, annealing 60°C for 10 seconds, and extension 72°C for 1 minute. Fragments were circularized using the NEBuilder reaction mix. Plasmids containing the donor sequence for recombination were constructed by digesting pGem-Easy with BamHI and HindIII restriction enzymes. PCR fragments were generated and circularized as described above. All plasmids were confirmed to have the intended sequences by Sanger sequencing.

### Protein Expression and Purification

Protein expression was performed in BL21 (DE3) strain of *E. coli*. Transformants were grown at 37°C in Luria-Bertani high salt (LB) media (supplemented with 100ug/ml ampicillin) with vigorous shaking until the OD_600_ reached ∼0.6. Protein expression was induced by adding isopropyl β-D-1-thiogalactopyranoside (IPTG) to a final concentration of 1mM and continued overnight at 18°C. The cells were pelleted by centrifugation at 8,000g for 20 minutes at 4°C. The pellet was suspended in lysis buffer (20mM Tris-Cl pH 8.0, 5mM imidazole, 200mM NaCl) and disrupted by sonication on ice. The insoluble fraction was removed by centrifugation at 15,000g for 15 minutes at 4°C. The supernatant was loaded onto a Ni^2+^-precharged 5ml GE HisTrap FF Crude column (GE Healthcare Life Science, Chicago, IL, United States). Non-specifically bound material was removed by a low imidazole wash buffer (20mM Tris-Cl pH 8.0, 50mM imidazole, 200mM NaCl) and bound material was collected using a high imidazole elution buffer (20mM Tris-Cl pH 8.0, 200mM imidazole, 200mM NaCl). Protein-containing fractions were then dialyzed overnight at 4°C in 2L of 50mM Tris-HCl pH 8.0. The dialyzed protein had the His-tag removed by the TEV protease with a TEV:protein ratio of 1:10 and with the addition of DTT and EDTA to final concentrations of 3 mM and 1.5 mM, respectively, to the reaction mixture. The digestion was carried over with simultaneous dialysis into 10mM Tris-HCl pH 8.0 buffer overnight. Protein(s) were further purified by ion-exchange chromatography using a GE MonoQ 10/100 GL column and 0 to 650 mM NaCl gradient in a 20 mM Tris-HCl pH 8.0 buffer. The fractions containing the target protein were pooled together and passed through a Ni^2+^-precharged 5ml GE HisTrap FF Crude column to remove the undigested protein. The final purification step was size exclusion chromatography using a Superdex 200 HiLoad 16/60 column in a buffer containing 20mM Tris-HCl pH 8.0, and 150mM NaCl. Proteins were concentrated and stored at 4°C.

### Circular Dichroism Assay

Full length gpV proteins were concentrated to 0.25mg/ml and dialyzed into 10mM NaPO4 pH 7.0 buffer overnight. The CD spectra were measured in the wavelength range of 190-250 nm at a temperature of 20°C using a JASCO J-815 CD Spectrometer. The final spectrum curves were the average of three spectra for each sample.

### Differential Scanning Fluorimetry Assay

DSF was conducted as described [23]. The proteins were concentrated to 1 mg/ml in 20mM Tris-HCl pH 8.0, and 150mM NaCl and mixed with an equal volume of SYPRO Orange dye, which was diluted 500 times from the stock concentration of 5000× (as supplied and labeled by ThermoFisher). The DSF experiments were carried out with a StepOnePlus™ Real-Time PCR System (Thermo Fisher Scientific). The temperature was continuously increased from 20°C to 99°C at a rate of 1.0 °C/min. Melting curves were directly exported from the instrument and then were analyzed with Thermo Fisher Software v1.3. Six melting curves were recorded for each protein.

### Crystallization, Data Collection, and Structure Determination

For crystallization, gpVdL variants were concentrated to 50mg/ml. All three mutants – gpVdL-AA, gpVdL-FF, gpV-DEL – crystallized in conditions similar to those found for gpVdL with the WT sequence: 26-33% Pentaerythritol ethoxylate (15/4 EO/OH) aka PEE-797, 100mM Bis-Tris or MES pH 6.0-6.5. The mother liquor served as a cryoprotectant for flash freezing in liquid nitrogen. Data collection strategy optimized the quality and completeness of high resolution bin (**Table 1**).

The X-ray diffraction data were integrated, reduced, and scaled using XDS [38, 39]. The crystal structures were solved by the molecular replacement method using Phaser [40, 41]. Various CCP4 programs were used in data manipulations [42]. The crystallographic refinement was performed using Refmac5 [43] and Phenix [44]. The atomic models were built and refined using Coot [45] (**Table 1**).

### Mutagenesis of Bacteriophage P2

Non-suppressing *E. coli* strain C-2 (the original P2 strain [26]) was transformed with the plasmid pACBSR, which contained the λ red system [46], and pGEM-T Easy containing a gpVam donor sequence and plated onto agar plates containing 100µg/ml ampicillin and 11µg/ml chloramphenicol (**Fig. 4**). After overnight incubation, multiple colonies were used to inoculate 1ml of LB, which was supplemented with 100µg/ml ampicillin and 11µg/ml chloramphenicol, to reach an OD_600_ of ∼0.1. Arabinose was added to a final concentration of 0.2% w/v and the culture was incubated for 1 hour at 37°C with vigorous shaking. Then, the culture was infected with a total of ∼10^6^ plaque-forming units (pfu) of P2 vir1 and, at the same time, CaCl_2_ was added to a final concentration of 5mM. After a 10 minute incubation with vigorous shaking, the cells were pelleted by centrifugation at 4,000g for 10 minutes at room temperature and gently resuspended in fresh media containing the antibiotics and arabinose. This removed the unabsorbed phage. After 25 minutes, the culture was supplemented with EDTA pH 8.0 at a final concentration of 0.8mM to prevent readsorption of the progeny phage. The infection was allowed to continue for 1 hour. The cell debris and non-lysed cells were pelleted by centrifugation at 5,000g for 10 minutes at room temperature.

The supernatant, which contained the progeny phage – a mixture of WT and *Vam*, was 10-fold serially diluted and plated on amber suppressing *E. coli* strain C-520 using dual-layer agar assay (the top layer contained 5mM CaCl_2_) to obtain well-isolated plaques. Then, ∼500 individual plaques were spot-tested on *E. coli* C-2 and C-520, and three mutants containing an amber mutation were identified. The material from these plaques was resuspended in 500µL of SM Buffer (50mM Tris-HCl pH 7.5, 100mM NaCl, 8mM MgSO_4_) and amplified by PCR. Sanger sequencing confirmed the presence of the designed *Am* mutation in the *V* gene of these mutants.

The resulting P2 vir1 *Vam* phage was used in the same recombination procedure with four different donor sequences – WT gpV, gpV-AA, gpV-FF, and gpV-DEL – to obtain P2 vir1 *Vam1-WT*, P2 vir1 *Vam1-AA*, P2 vir1 *Vam1-FF*, and P2 vir1 *Vam1-DEL* phage mutants. The post-recombination selection procedure was performed on non-amber suppressing *E. coli* C-2. It required only a single step because P2 vir1 *Vam1* did not propagate on this strain. The presence of designed mutations was confirmed by Sanger sequencing.

### Purification of Bacteriophage P2

For each 1L of the final culture, 100 ml of *E. coli* C-2 was first grown in the LB medium to OD_600_ of ∼1.0 at 37°C with vigorous shaking. 10ml of this seed culture were supplemented with 1mM CaCl_2_ and infected with ∼1×10^6^ pfu of P2 while the 90 ml fraction was set to rest at room temperature. The infected fraction was incubated for 10 minutes at 37°C in a heat block. It was then mixed with the 90ml fraction and the mixture was added to 1 L LB medium, which was supplemented with 0.5mM CaCl_2_, 1.6mM MgCl_2_, and 0.1% Glucose. The culture was then incubated at 37°C with vigorous shaking and its OD_600_ was recorded every 10 minutes. When a reduction in OD_600_ was detected (usually at OD_600_ of ∼0.7-1.5), which indicated the beginning of lysis, EDTA pH 8.0 was added to a final concentration of 0.3% w/v to prevent the readsorption of progeny phage. Once the culture had cleared (its OD_600_ dropped to ∼0.1), cellular debris was removed by centrifugation at 5,000g for 20 minutes at +4°C. Phages were precipitated by the addition of PEG 6,000 (to 8% w/v) and NaCl (to reach 2.5% w/v, taking into account 1% NaCl in the LB) overnight at 4°C. The pellet was resuspended in 10ml SM buffer and excess lipids were removed by centrifugation at 15,000g for 15 minutes at 4°C, and the phage was concentrated by centrifugation at 100,000g for 1.5 hours at 4°C. The supernatant was decanted and the pellet, containing the phage, was resuspended in 0.5ml SM buffer overnight at +4°C with moderate shaking. The sample was further purified by isopycnic ultracentrifugation in a CsCl gradient. The phage-containing sample was layered on top of a CsCl solution in SM buffer with a density of 1.45mg/ml and centrifugated overnight at 100,000g at +4°C. The phage-containing opalescent band was extracted from the gradient with a syringe and dialyzed into SM buffer with 3 changes of buffer at two 2h time points and then overnight.

### EOP Assay

The purity of phage samples was evaluated using SDS-PAGE stained with a Coomassie Brilliant Blue dye. The relative abundance of fully assembled phage particles was derived from the intensities of the gpFI sheath and gpFII tube protein bands that were determined using ImageJ software [47]. Phage (WT and mutants) titers were first established using 10 fold serial dilution dual-layer agar assay in which the phage sample was drop-spotted on *E. coli* C-2 in soft agar supplemented with 5mM CaCl_2_. The equivalent ∼100 pfu of phage was added to ∼1×10^8^ colony forming units (cfu) of *E. coli* C-2, incubated overnight at 37°C, and individual plaques were counted.

### Central Spike Localization Assay and Western Blot

Localization was conducted as described previously (Mattenberger et al.) with a small modification – rifampicin was added to stop the transcription of phage genes. Briefly, *E. coli* C-2 was grown in LB media and supplemented with 25 μg/mL rifampicin 10 minutes prior to phage infection. 100 mL of LB culture containing *E. coli* C-2 with an OD_600_ of 0.1 were mixed with P2 (WT and mutants) at an MOI of 20 (2×10^11^ pfu of phage) or an equal volume of SM Buffer. CaCl_2_ was added to a final concentration of 5 mM. The mixture was incubated with shaking at 37°C for 15 minutes and then centrifugated at 5,000g for 5 minutes. The supernatant was saved to determine the efficiency of absorption. The pellet was resuspended in ice-cold Tris-HCl pH 7.5 buffer and pelleted again by centrifugation at 5,000g for 5 minutes. The cell pellet was suspended in 1 ml of sucrose-EDTA-lysozyme buffer (200 mM Tris-HCl, pH 8.0, 500 mM sucrose, 1 mM EDTA, 1 mg/mL lysozyme) and incubated in an ice water bath for 30 minutes and then centrifugated at 16,000g for 30 minutes. The supernatant, which contained the periplasmic fraction, was collected.

To separate the membrane and cytoplasmic fractions, the pellet was resuspended in 10 ml of ice-cold Tris-HCl, pH 7.5 and sonicated. Large insoluble components were removed by centrifugation at 5,000g for 5 minutes. To isolate the membranes, the supernatant was centrifugated at 100,000g for 1.5 hours. The resulting pellet contained both membranes while the supernatant contained the cytoplasmic fraction.

Western blot procedure was conducted as previously described using the same set of markers (Mattenberger et al.). The primary antibodies were as follows: MBP-probe antibody (Santa Cruz Biotechnology, sc-13564), GroEL polyclonal antibody (Enzo Life Sciences, ADI-SPS-875-D), OmpF antibody (orb308741, Biorbyt) and a custom made anti-P2 gpV polyclonal antibody generated by GenScript. The secondary antibodies were both from Abcam: a goat polyclonal antibody to rabbit (ab6721) and a goat polyclonal antibody to mouse (ab6789). Phages used in the localization assay were titrated on *E. coli* C-2 lawn prior to and post host cell adsorption to evaluate the efficiency of absorption. In the case of post-adsorption cell culture, the infected cells were pelleted down and the titer of the resulting supernatant was assayed.

### Molecular Graphics

Figures 1c and 4a were created using UCSF Chimera [48]. Figure 8 was made using UCSF ChimeraX [49].

## Author contributions

**P.G.L**. and **J.M.M**. conceived the study. **J.M.M**. performed mutagenesis, cloned, purified, crystallized, and conducted CD and DSF experiments of WT gpV and mutant proteins. **J.M.M**. performed mutagenesis, purified phage and conducted infectivity assays. **E.S.K**. performed cellular localization assay and western blots. **S.A.B**. and **N.S.P**. assisted with the initial development of protein purification protocols and phage mutagenesis, respectively. **P.G.L**. collected and processed the X-ray crystallography data. **J.M.M**. and **P.G.L**. wrote the manuscript, which was read, edited, and approved by all authors.

## Correspondence and requests

Correspondences should be addressed to Petr G. Leiman.

## Acknowledgments

This work was supported by the UTMB Sealy Center for Structural Biology and Molecular Biophysics, by the Department of Biochemistry and Molecular Biology, and by the National Institute of Health (Grant R01GM139034). We thank Dr. Andres Oberhauser for use of his RT-PCR machine, and Dr. Luis Holthauzen for his help in CD and DSF analysis. We also thank Richard Calendar for providing P2 vir1 and *E. coli* strains used in this study. X-ray diffraction data were collected at the LS-CAT Sector 21 beamlines of the Advanced Photon Source. We acknowledge the use of the Advanced Photon Source, a U.S. Department of Energy (DOE) Office of Science User Facility operated for the DOE Office of Science by Argonne National Laboratory under Contract No. DE-AC02-06CH11357. We thank the staff of the LS-CAT Sector 21 beamlines that is supported by the Michigan Economic Development Corporation and the Michigan Technology Tri-Corridor (Grant 085P1000817).

## Supplementary Information

is available for this paper.

