## Supplementary Table 1 for "Function of the bacteriophage P2 baseplate central spike Apex domain in the infection process"

**Inventory of Supplemental Information**

**Supplementary Tables**

- **Supplementary Table 1 List of primers for cloning of gpV mutants and gpVdL mutants.**

**Supplementary Table 1. List of primers for cloning of gpV mutants and gpVdL mutants.**

Primers used for gpV cloning and mutagenesis (capitalized is Bacteriophage P2 sequence, highlighted nucleotides are mutated).

| **Mutation** | **Primer** | **Sequence** |
| --- | --- | --- |
| HxH->AxA | gpV-AxA-F | CTGTCGCCGGGCGCTTTCGCGGTATGCAGTACCTTACCGTTTGATGAGAG |
|  | gpV-AxA-R | AAGGTACTGCATACCGCGAAAGCGCCCGGCGACAGCGGCGGCACA |
| HxH->FxF | gpV-FxF-F | CTGTCGCCGGGGAATTTGAAGGTATGCAGTACCTTACCGTTTGATGAGAG |
|  | gpV-FxF-R | AAGGTACTGCATACCTTCAAATTCCCCGGCGACAGCGGCGGCACA |
| HxH->DEL | gpV-DEL-F | GCCGGGAAATTTAAagggtatgcagtacctt |
|  | gpV-DEL-R | tttaaATTTCCCGGCGACAGCGGCGGCACA |
| Vam Universal Recombination Forward | gpVam-URF | ggcggccgcgggaattcgatGAGGCTGATGCGCAATTAAACAT |
| Vam  Recombination Forward | gpVam-RF | ACTCTCGCAAATATTTAGGAACTCG |
| Vam  Recombination Reverse | gpVam-RR | ATTTGCGAGAGTGTTCATGCATGT |
| Universal Recombination Reverse | gpV-URR | gccgcgaattcactagtgatcgaaTCACTCATCCGAGCCTCC |
| HxH->DEL Recombination | gpV-DEL-RF | GCATACCTTATGACAGCGCGTTATCTCG |
|  | gpV-DEL-RR | CGCTGTCATAAGGTATGCAGTACCTTACCGTT |
